# HVEM structures and mutants reveal distinct functions of binding to LIGHT and BTLA/CD160

**DOI:** 10.1101/2021.06.16.448617

**Authors:** Weifeng Liu, Ting-Fang Chou, Sarah C. Garrett-Thomson, Goo-Young Seo, Elena Fedorov, Udupi A. Ramagopal, Jeffrey B. Bonanno, Kiyokazu Kakugawa, Hilde Cheroutre, Mitchell Kronenberg, Steven C. Almo

## Abstract

HVEM is a TNF (tumor necrosis factor) receptor contributing to a broad range of immune functions involving diverse cell types. It interacts with a TNF ligand, LIGHT, and immunoglobulin (Ig) superfamily members BTLA and CD160. Assessing the functional impact of HVEM binding to specific ligands in different settings has been complicated by the multiple interactions of HVEM and HVEM binding partners. To dissect the molecular basis for multiple functions, we determined crystal structures that reveal the distinct HVEM surfaces that engage LIGHT or BTLA/CD160, including the human HVEM:LIGHT:CD160 ternary complex, with HVEM interacting simultaneously with both binding partners. Based on these structures, we generated mouse HVEM mutants that selectively recognized either the TNF or Ig ligands *in vitro*. Knock-in mice expressing these muteins maintain expression of all the proteins in the HVEM network, yet they demonstrate selective functions for LIGHT in the clearance of bacteria in the intestine and for the Ig ligands in the amelioration of liver inflammation.

## Introduction

Members of the tumor necrosis factor receptor super family (TNFRSF) regulate diverse processes, but in several cases understanding these processes is hampered by the ability of receptors and ligands to bind to multiple partners (Bossen et al., 2006). One prominent example is provided by the herpes virus entry mediator (HVEM), or TNFRSF14, initially identified as important for entry of herpes simplex virus (HSV) through recognition of HSV glycoprotein D (gD) (Montgomery et al., 1996; Whitbeck et al., 1997). Subsequently, a TNF super family (TNFSF) ligand for HVEM was characterized, known as **LIGHT** (homologous to **l**ymphotoxin, exhibits **i**nducible expression and competes with HSV **<u>g</u>**lycoprotein D for binding to **<u>h</u>**erpesvirus entry mediator, a receptor expressed on **<u>T</u>** lymphocytes) or TNFSF14 (Harrop et al., 1998a; Harrop et al., 1998b). Engagement of HVEM by LIGHT is implicated in multiple responses. For example, in T lymphocytes, it stimulates proliferation, cytokine production, and the development of CD8 T cell memory (Desai et al., 2017; Harrop et al., 1998a; Harrop et al., 1998b; Tamada et al., 2000). LIGHT also engages HVEM to stimulate cytokine production by type 3 innate lymphoid cells (ILC3) (Seo et al., 2018) and in keratinocytes it binds HVEM to stimulate periostin, contributing to atopic dermatitis (Herro et al., 2018).

LIGHT also binds to another TNFRSF member, lymphotoxin-beta receptor (LT*β*R or TNFRSF3), which is expressed by stromal and myeloid lineages. This interaction regulates lymph node formation, dendritic cell migration (Zhu et al., 2011), and IL-12 production by DC (Okwor et al., 2015). The LIGHT-LT*β*R interaction also has been reported to induce apoptosis of cancer cells (Zhai et al., 1998), it is important for macrophage activity in wound healing (Petreaca et al., 2012) and it influences lipid metabolism by regulating hepatic lipase expression in hepatocytes (Chellan et al., 2013; Lo et al., 2007). Furthermore, LIGHT participates in additional processes in which a specific receptor has not been implicated, including the resolution of inflammation in an experimental autoimmune encephalomyelitis (Mana et al., 2013), the induction of adipocyte differentiation (Tiller et al., 2011), and the induction of osteoclastogenic signals (Brunetti et al., 2014; Hemingway et al., 2013).

HVEM also binds immunoglobulin superfamily (IgSF) molecules B and T lymphocyte attenuator (BTLA or CD272) and CD160. HVEM engages in bidirectional signaling, serving not only as a receptor, but it also may act as a ligand for IgSF receptor signaling (Steinberg et al., 2011). HVEM:BTLA engagement delivers an overall inhibitory immune response (Murphy and Murphy, 2010), while the interaction between HVEM and CD160 on T cells can either attenuate the activities of specific subsets of CD4 T lymphocytes or enhance the activity of CD8 T cells (Cai et al., 2008; Tan et al., 2018). Notably, engagement of CD160 by HVEM also controls cytokine production by NK cells and is important for mucosal immunity (Shui et al., 2012; Tu et al., 2015; Whitbeck et al., 1997). Furthermore, HVEM was reported to interact with synaptic adhesion-like molecule 5 (SALM5), mainly expressed in brain, to confer immune-privilege in the central nervous system (Zhu et al., 2016). These different interactions are summarized in **Fig. S1**. CD160 also binds to some major histocompatibility complex (MHC) class I molecules (Le Bouteiller et al., 2002; Maeda et al., 2005), further expanding the complexity of this protein-protein interaction network.

The promiscuous interactions of HVEM pose challenges for characterizing the mechanistic contributions of HVEM-associated pathways in different immune responses and diseases. Conditional knockouts can isolate effects in particular cell types, but elimination of expression of one protein, for example LIGHT, not only abolishes LIGHT-HVEM binding, but also eliminates LIGHT-LT*β*R binding and may also indirectly affect HVEM interactions with its IgSF ligands by altering the availability of HVEM (Steinberg et al., 2011). This complexity may make it difficult to reach definitive conclusions about the relevant binding partners responsible for a phenotype and it may account for circumstances in which the phenotypes in whole body receptor and corresponding ligand knockouts did not agree (Giles et al., 2018). Herein, in order to better understand this receptor-ligand network, we set out to test mutants of HVEM with selective ligand binding. Based on multiple crystal structures, including the human HVEM:LIGHT:CD160 ternary complex we performed extensive epitope mapping and engineering of selective mHVEM mutants. HVEM muteins were expressed in mice to show definitively that selective HVEM- ligand interactions are important in resistance to mucosal bacterial infection and in prevention of liver inflammation in a context where all members of the protein network were present and only selective interactions were disrupted.

## RESULTS

### Human HVEM:LIGHT complex exists as a 3:3 assembly

The extracellular domains of human LIGHT (denoted as hLIGHT; ∼18 KDa for the monomer and ∼54 KDa for the homotrimer) and human HVEM (denoted as hHVEM; ∼15 KDa) were purified to homogeneous, monodisperse species as indicated by analytical size exclusion chromatography (SEC) **(Fig. 1 A)**. Mixing equal molar equivalents of hLIGHT and hHVEM monomers, resulted in a single species with an apparent molecular weight of ∼100 KDa, consistent with the formation of a 3:3 stoichiometric hHVEM:hLIGHT assembly in solution (**Fig. 1, A-C**).

**Figure 1.**
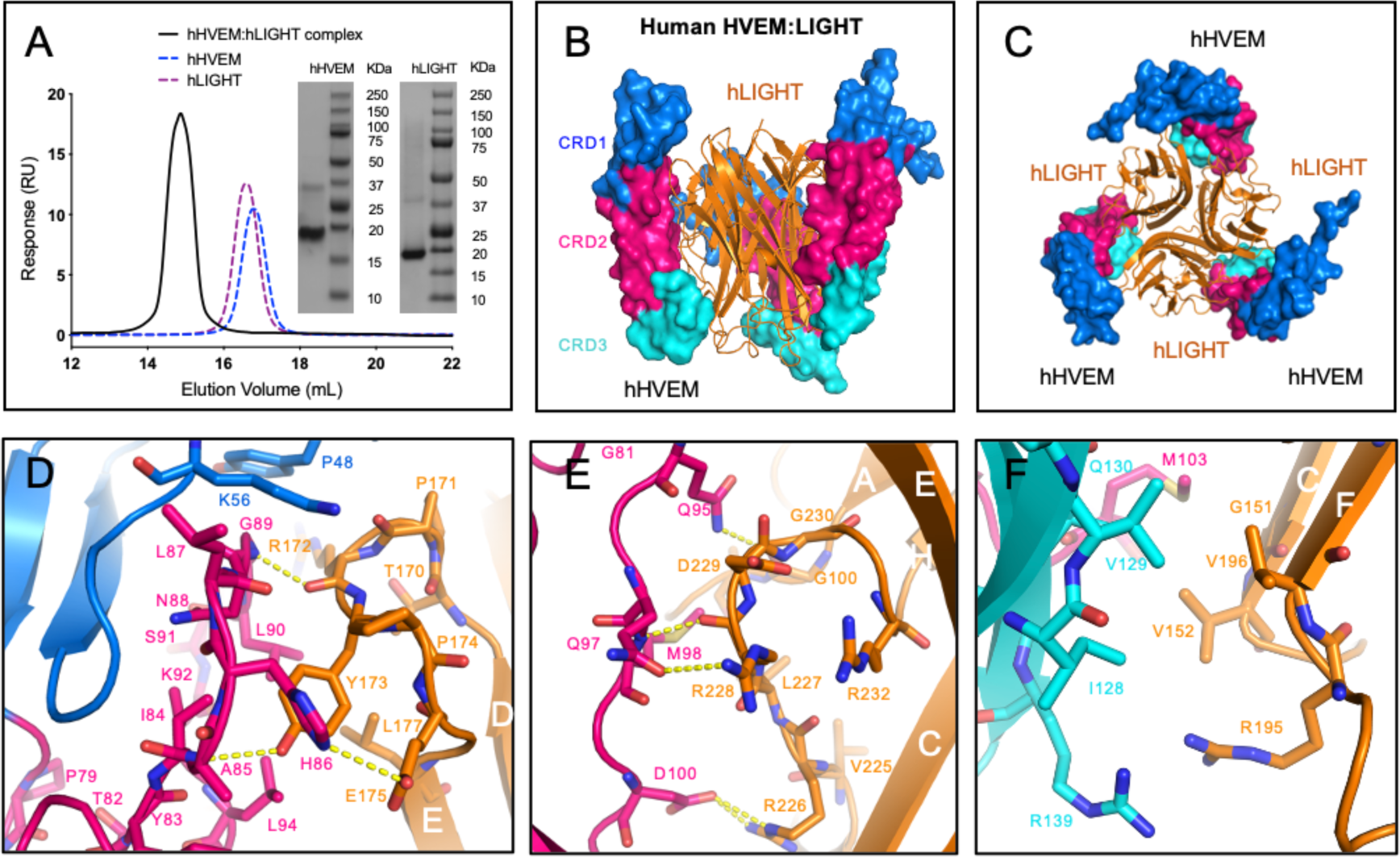
Crystal structure of human HVEM:LIGHT complex exhibits a 3:3 stoichiometry. **(A)** The analytical SEC trace of hHVEM and hLIGHT mixtures reveals a significant peak of the complex corresponding to the molecular weight around 100 kDa. The SDS-PAGE results indicate hHVEM and hLIGHT were purified to near homogeneity. Note that in the SDS gel, LIGHT trimers dissociate. **(B and C)** The hHVEM is shown as a surface and each CRD domain is colored separately as indicated in the figure. The trimeric hLIGHT protein is shown as an orange ribbon in the figure. The side view **(B)** and bottom view **(C)** of the hHVEM:hLIGHT complex are shown. **(D-F)** The detailed interaction interface between hHVEM and hLIGHT. The hHVEM CRD1, CRD2, and CRD3 residues are colored as marine, hot pink, and cyan, respectively. hLIGHT residues are colored as orange. The hydrogen bonds between hHVEM and hLIGHT are indicated as dashed lines.

The crystal structure of the hHVEM:hLIGHT complex was determined to the resolution of 2.30 Å by molecular replacement using Protein Data Bank (PDB) entries 4KG8 (hLIGHT) and 4FHQ (hHVEM) as starting search models (**Table 1**). The asymmetric unit of the hHVEM:hLIGHT crystals contains six independent chains of hLIGHT and six independent chains of hHVEM, which form two classical 3:3 TNF:TNFR hexameric assemblies with three-fold symmetry (**Fig. S2, A-C**); a single 3:3 TNF:TNFR hexameric assembly is consistent with SEC analysis. The hHVEM ectodomain is composed of four cysteine rich domains (CRDs), while hLIGHT forms a compact homotrimeric structure. In the hexameric assembly, CRD1, CRD2 and CRD3 of hHVEM engage hLIGHT via surfaces contributed by two adjacent hLIGHT protomers (**Fig. 1 B and S2 C**). The two independent hHVEM:hLIGHT hexameric complexes exhibit similar overall structures with a RMSD of 1.8 Å for 742 aligned C*α* atoms. The regions with the greatest structural divergence reside in the N- and C-termini of the proteins, which do not directly contribute to the binding interface. The hHVEM:hLIGHT recognition interfaces are highly similar within and between the two complexes (**Fig. S2 B**), and the following discussion is based on the hLIGHT G and H chains, and hHVEM J chain (**Fig. S2 A**).

**Table 1.** Data Collection and Refinement Statistics

|  | hHVEM:hLIGHT | hHVEM:hLIGHT:hCD160 | mHVEM |
| --- | --- | --- | --- |
| **Data Collection** |  |  |  |
| Wavelength used (Å) | 1.075 | 0.97931 | 0.97931 |
| Resolution range (Å) | 2.30-50.00<br>(2.30-2.34) | 3.50-50.00<br>(3.50-3.83) | 2.10-50.00<br>(2.10-2.14) |
| Space group | P2<sub>1</sub>2<sub>1</sub>2<sub>1</sub> | I23 | P4<sub>1</sub>2<sub>1</sub>2 |
| Unit cell (Å) | a=111.7,<br>b=113.6,<br>c=163.3 | a=b=c=214.7 | a=b=64.7,<br>c=69.0 |
| Unique reflections (N) | 92792 | 20868 | 8989 |
| Redundancy | 10.8(10.7) | 20.7 (17.9) | 13.5 (9.9) |
| Completeness | 99.9(99.7) | 99.9 (100) | 99.5 (99.1) |
| I/sigma | 22.7 (3.1) | 16.1 (2.2) | 17.1 (2.2) |
| R<sub>merge</sub> | 0.125 (0.936) | 0.191 (1.674) | 0.135 (0.938) |
| CC<sub>1/2</sub> | N/A | 0.999 (0.676) | 0.999 (0.943) |
| **Refinement** |  |  |  |
| Resolution range (Å) | 2.30-48.92<br>(2.30-2.36) | 3.50-19.93<br>(3.50-3.59) | 2.10-20.00<br>(2.10-2.16) |
| R<sub>work</sub> | 0.188 (0.245) | 0.257 (0.370) | 0.212 (0.355) |
| R<sub>free</sub> | 0.231 (0.270) | 0.285 (0.293) | 0.257 (0.328) |
| Average B factor (Å<sup>2</sup>) | 38.4 | 139.9 | 55.5 |
| Rms bond (Å) | 0.021 | 0.005 | 0.018 |
| Rms angles (°) | 2.081 | 1.290 | 1.928 |
| PDB code | 4RSU | 7MSG | 7MSJ |

Parentheses indicate statistics for the highest resolution bin.

### The binding interface between human HVEM and LIGHT

The structure of the hHVEM:hLIGHT complex shows that HVEM CRD1 and CRD2 domains interact with the DE, AA’ and GH loops of LIGHT, while HVEM CRD3 interacts with LIGHT CD and EF loops (**Fig. 1, D-F and S2, C-D**).

The interaction between the hHVEM CRD2 and the hLIGHT DE loop appears to be important for human HVEM:LIGHT recognition, as it contributes multiple potential polar contacts. The main chain amide group of hHVEM A85 forms a hydrogen bond with the side chain hydroxyl group of hLIGHT Y173 (**Fig. 1 D and S2 D**), consistent with the behavior of the Y173F mutation in hLIGHT, which significantly diminishes the binding of hLIGHT with hHVEM (Rooney et al., 2000). Human HVEM N88 does not directly contact hLIGHT Y173, but is relatively close, and the hHVEM N88A mutation attenuated binding to hLIGHT (**Fig. S3, A-D**). The hHVEM G89 main chain amide group forms a hydrogen bond with the main chain oxygen of hLIGHT R172 (**Fig. 1 D and S2 D**). HVEM H86 side chain imidazole functionality makes a polar contact with the side chain carboxyl group of hLIGHT E175 (**Fig. 1 D and S2 D**). It was previously reported that the hHVEM H86I mutation dramatically reduced binding to hLIGHT (Shrestha et al., 2020).

Human HVEM CRD2 forms four additional polar contacts with GH loop of hLIGHT (**Fig. 1 E and S2 D**). The hHVEM Q97 side chain oxygen forms a polar contact with hLIGHT R228 side chain. Human HVEM M98 backbone amide group contacts the backbone oxygen of hLIGHT R228 and the side chain carboxyl group of hHVEM D100 forms two polar contacts with the side-chain guanidinium group of hLIGHT R226 (**Fig. 1 E and S2 D**). The hHVEM D100R mutation resulted in undetectable binding with hLIGHT (Shrestha et al., 2020). The AA’ loop from the lower region of CRD2 contributes only a single polar contact, formed by the main chain oxygen from G100 of hLIGHT and the side chain amide group of hHVEM Q95 (**Fig. 1 E and S2 D**).

hHVEM CRD3 residues, including I128-G132, H134 and A136-R139 participate in interactions with G151-V152 and A159-T161 from the CD loop, as well as residues Q183, R195-V196 and W198 from the EF loop of hLIGHT (**Fig. S2, C and D**). Examination of the structure in this region reveals no polar contacts between hHVEM and hLIGHT. A modest hydrophobic interface is formed by the packing of the side chains of hHVEM residues I128 and V129 against the side chains of hLIGHT V152 and V196 (**Fig. 1 F and S2 D**).

### Structure of the human HVEM:LIGHT:CD160 ternary complex

It was previously shown that LIGHT and the IgSF ligands do not compete for binding to HVEM (Cai et al., 2008; Liu et al., 2019), suggesting the potential for forming a ternary complex. Therefore, we set out to solve the crystal structure of hHVEM:hLIGHT:hCD160 (human CD160 is denoted as hCD160) complex (PDB entry 7MSG). Accordingly, we determined the structure of this complex to 3.5 Å resolution by molecular replacement using CD160 (PDB entry 6NG9) and the hHVEM:hLIGHT complex described above (PDB entry 4RSU) as search models (**Fig. 2, A and B**). The asymmetric unit contains three copies of each hHVEM, hLIGHT and hCD160, forming a ternary complex with 3:3:3 stoichiometry. Within the ternary assembly, hHVEM and hLIGHT exhibit the classical 3:3 TNF:TNFR assembly, with contacts that are very similar to the structure of the hHVEM:hLIGHT binary complex described above. The hHVEM:hLIGHT complex forms the core of the ternary complex with each hHVEM CRD1 further binding a single molecules of hCD160 in a manner similar to that observed in the structure of the hHVEM:hCD160 binary complex (**Fig. 2, A and D and Fig. S3 C**). Notably, the structures of hHVEM:hLIGHT:hCD160 and hHVEM:hCD160 complexes relied on the use of a single chain hCD160-hHVEM fusion protein as the relatively weak interaction of hCD160-hHVEM (7.1 ± 0.9 μM) does not support the stable complex formation in solution (Liu et al., 2019). The crystal structure of the hHVEM:hLIGHT:hCD160 complex provides direct evidence that hLIGHT and hCD160 can simultaneously engage hHVEM, resulting in a higher order assembly with the potential of coordinated signaling through both hHVEM and hCD160. Notably, the simultaneous interaction of hCD160 and hLIGHT with hHVEM alters the local organization of hCD160, as engagement of hHVEM with trimeric hLIGHT may enforce close proximity of up to three hCD160 molecules with distinct geometric organization, as compared to the engagement of hCD160 and hHVEM in the absence of hLIGHT.

**Figure 2.**
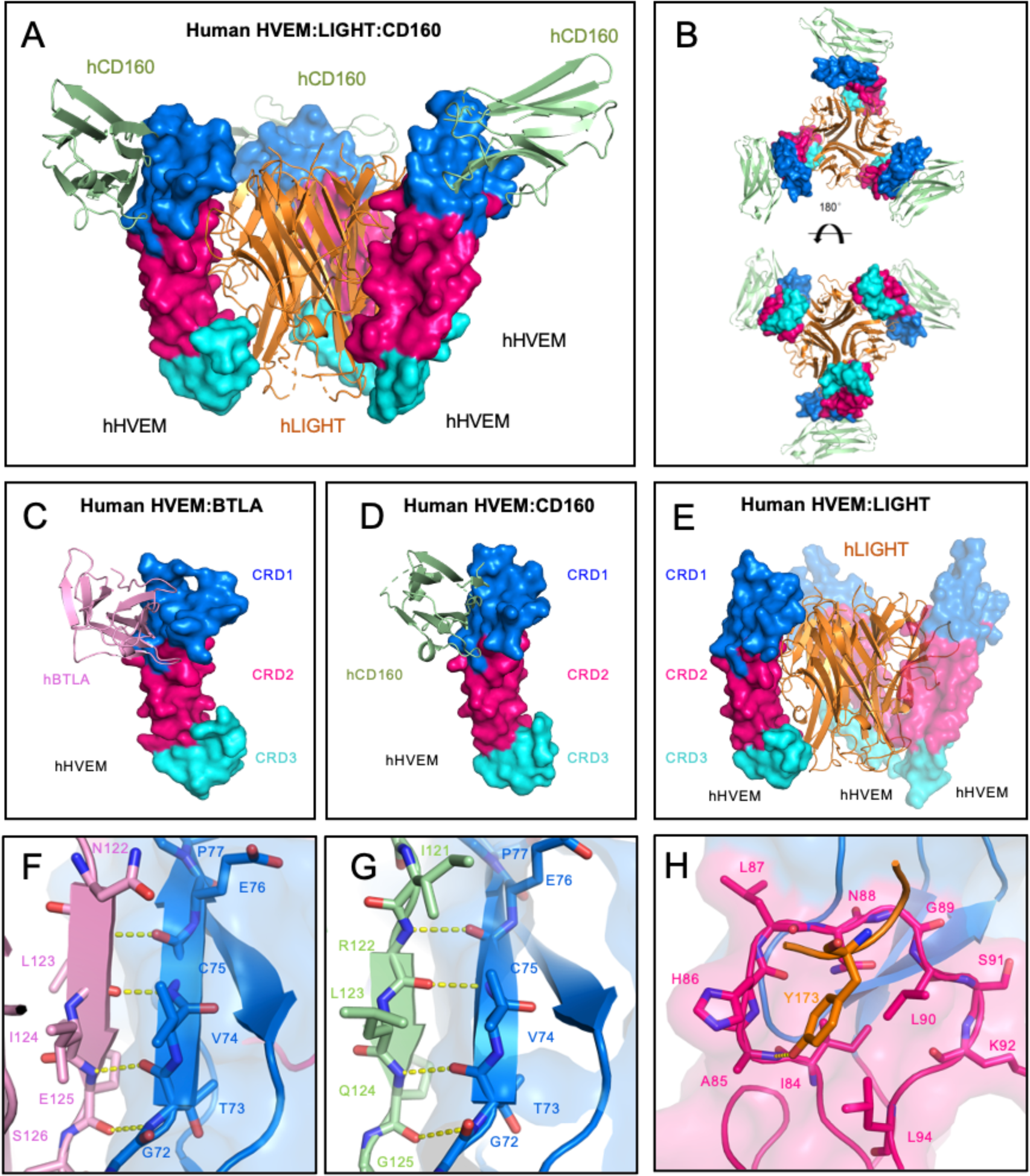
Overall structure of HVEM:LIGHT:CD160 ternary complex and critical interaction interfaces of HVEM binding to BTLA, CD160 and LIGHT. **(A and B)** Structure of the hHVEM:hLIGHT:hCD160 ternary complex indicates hCD160 and hLIGHT can interact simultaneously with hHVEM. The side view **(A)** and the top/bottom views **(B)** of the ternary complex are shown. **(C)** Structure of hHVEM:hBTLA (PDB entry 2AW2). **(D)** Structure of hHVEM:hCD160 (PDB entry 6NG3). **(E)** Structure of hHVEM:hLIGHT (PDB entry 4RSU). These structures indicate hBTLA and hCD160 bind to similar surfaces on hHVEM, whereas hLIGHT binds to a different surface on hHVEM. **(F-H)** Detail binding interfaces between hHVEM and its binding ligands hBTLA, hCD160 and hLIGHT, respectively. The hHVEM CRD1 and CRD2 domains are colored as marine and hot pink, respectively.

Crystal structures and complementary mutagenesis studies of hHVEM:hCD160 and hHVEM:hBTLA (human BTLA is denoted as hBTLA) complexes demonstrated that both hCD160 and hBTLA mainly bind to CRD1 on hHVEM (**Fig. 2, C and D**) (Compaan et al., 2005; Liu et al., 2019). In contrast, the crystal structure of the hHVEM:hLIGHT complex shows hLIGHT binds to CRD2, CRD3 and a small part of CRD1 on hHVEM (**Fig. 2 E**). Crystal structures of hHVEM in complex with hBTLA and hCD160 highlight an anti- parallel intermolecular *β−*strand interaction, in which the *β−*strand composed of residues G72-P77 from CRD1 in hHVEM contacts the edge *β−*strands in hBTLA and hCD160 through canonical main-chain-to-main-chain *β−*sheet hydrogen bonds (**Fig. 2, F and G**). This pattern of hCD160 interactions with hHVEM is conserved in the ternary hHVEM:hLIGHT:hCD160 complex. Mutations of residues within this intermolecular *β−*strand (G72-P77) in HVEM CRD1 significantly altered the binding affinities (Shrestha et al., 2020), while hHVEM CRD2 mutations do not significantly alter the affinities to hCD160 and hBTLA. In contrast, HVEM CRD2 mutations, particularly the HVEM residues forming the concave cavity surrounding hLIGHT Y173, significantly affect hHVEM:hLIGHT binding (**Fig. 2 H**). Because both hCD160 and hBTLA bind to similar epitopes on hHVEM CRD1 (Compaan et al., 2005; Liu et al., 2019), it is likely that hHVEM, hLIGHT and hBTLA are able to form a ternary complex similar to the trimolecular complex of hHVEM:hLIGHT:hCD160 we have determined.

### Structure guided mutagenesis of mouse HVEM mutants

The mHVEM (mouse HVEM is denoted as mHVEM, PDB entry 7MSJ) structure was determined to 2.10 Å resolution by molecular replacement using the human HVEM (PDB entry 4FHQ) as the search model. The mouse and human HVEM structures are similar with RMSD of 2.7 Å for 97 aligned C*α* atoms, with the biggest differences in CRD3 (**Fig. 3, A and B**). Based on structural and sequence alignments between hHVEM and mHVEM, the solvent accessible mHVEM residues close to the putative binding interfaces were mutated to dissect the interaction network and enable in vivo HVEM functional studies.

**Figure 3.**
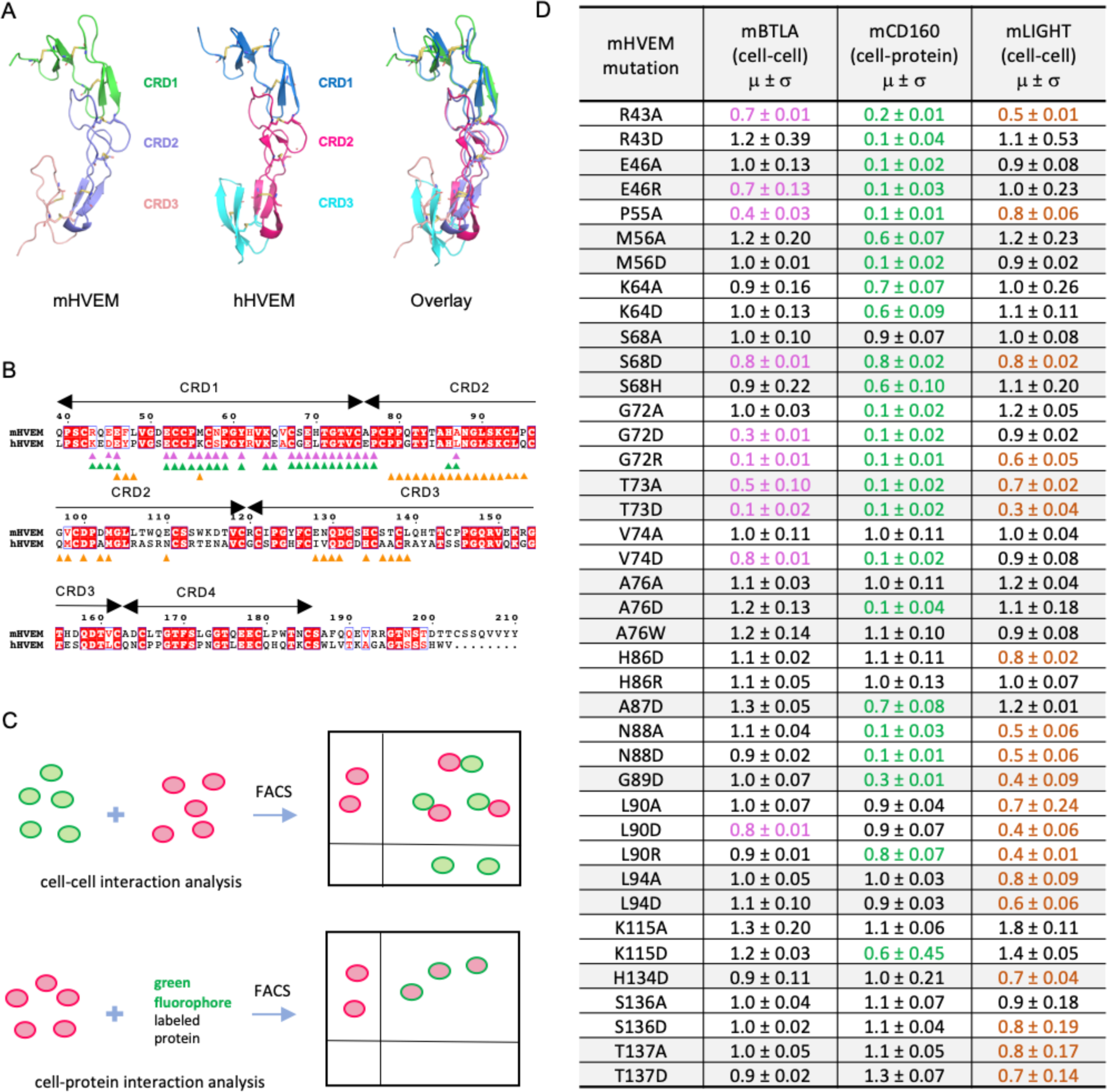
Structure and mutagenesis screen of mHVEM. **(A)** Structures of mHVEM, hHVEM and their comparison. The disulfide bonds of HVEM are shown as sticks and each HVEM CRD is colored differently. **(B)** Sequence alignment of mHVEM and hHVEM. The homologous residues are highlighted in red. The residues of hHVEM directly involved in the interface with hBTLA, hCD160, and hLIGHT are marked by magenta, green and orange triangles, respectively. **(C)** The schematic figure shows two ways to determine the relative binding affinities of mHVEM mutants. The cell-cell method measures the percentages of double positive cells in the mixtures. The cell-protein method measures the percentages of green-fluorophore stained mHVEM-mCherry expressing cells. **(D)** Relative binding affinities of mHVEM mutants with its ligands are shown in the table. Both mBTLA and mLIGHT binding to mHVEM was assessed by cell- cell method. The mCD160 binding to mHVEM was tested by cell-protein method. Error bars represent results from at least triplicates. All mHVEM mutants with ≥ 20% binding reduction to a particular query are colored differently to indicate their reduced affinities.

The relative binding affinities of mHVEM mutants with mBTLA and mLIGHT (mouse BTAL and LIGHT are denoted as mBTLA and mLIGHT, respectively) were evaluated by a cell-cell interaction assay (**Fig. 3 C**). The relative binding affinities of mHVEM mutants for mCD160 (mouse CD160 is denoted as mCD160) binding were screened using a cell-soluble protein assay because of low surface expression of the CD160 protein. A total of 52 mHVEM surface residues within or close to the likely ligand binding interfaces were individually mutated to different amino acids to probe the effect on ligand binding and to identify variants with selective ligand recognition (**Fig. 3 D**). For example, alteration of mHVEM G72 or V74 to aspartic acid attenuated binding to both mBTLA and mCD160, but not binding to mLIGHT; the mHVEM R43D, M56D or A76D mutations decreased binding to mCD160 but not mBTLA and mLIGHT; the mHVEM H86D, L90A, L94A and L94D mutations compromised the interaction with mLIGHT but not to mBTLA or mCD160 (**Fig. 3 D and S3 E**).

To further modulate the selectivity toward mLIGHT or mBTLA/mCD160, mHVEM mutations with similar binding properties were combined (**Fig. 4 A and S3 F**). For example, the combination of the G72 and V74 mutations completely eliminated binding to both mBTLA and mCD160, but did not appreciably impact mLIGHT binding in the flow cytometry based binding assays. Various pairwise combinations of mutations of H86, L90 and L94 eliminated mLIGHT binding, but did not substantially impact binding to mBTLA or mCD160 (**Fig. 4 A and S3 F-G**). Thus, these compound mutations resulted in several additional mHVEM variants with considerable binding selectivity. Although triple mutation of H86, L90 and L94 removed mLIGHT binding, it also dramatically reduced binding to mBTLA and mCD160 (**Fig. S3 F**). Not surprisingly, other combinations of mutations also reduced the binding to all ligands, such as the mHVEM R43D-M56A-K64D triple mutation (**Fig. S3 F**).

**Figure 4.**
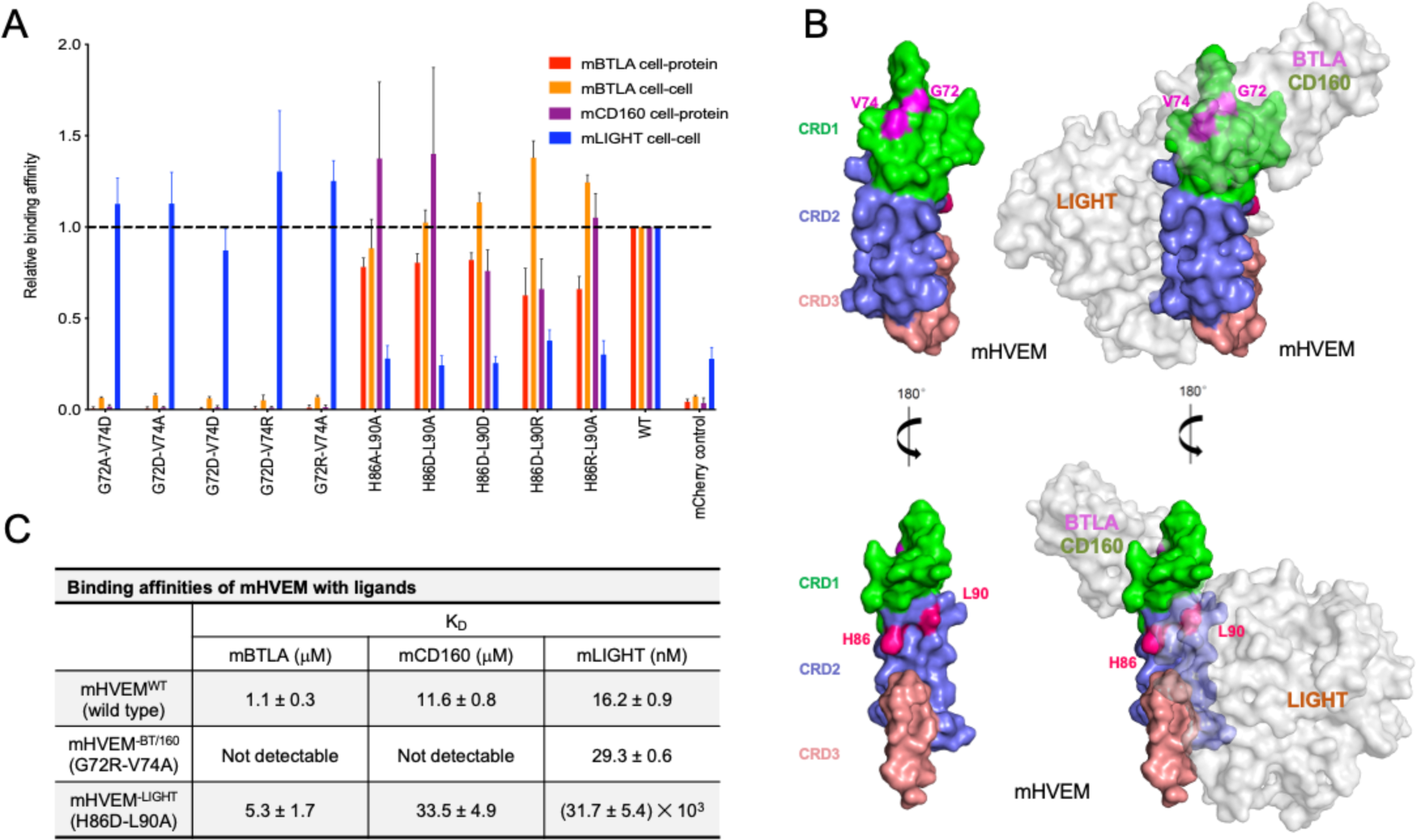
The engineered mHVEM mutants have binding selectivity. **(A)** The relative binding affinities of mHVEM mutants with mBTLA, mCD160, and mLIGHT as measured by cell-cell or cell-protein methods. Error bars represent results from at least triplicates. The grey dashed line marks the averaged normalized affinities of wild-type mHVEM with mBTLA, mCD160, and mLIGHT. **(B)** The locations of the mutated residues on mHVEM. mHVEM is shown as surface with each CRD colored differently with G72, V74, H86 and L90 are marked on the mHVEM surface. Ligands BTLA, CD160 and LIGHT are modeled based on the HVEM structures and are shown as labeled grey surfaces. **(C)** The binding affinities of mHVEM^WT^ (wild-type mHVEM), mHVEM^-BT/160^ (mHVEM G72R- V74A double mutein) and mHVEM^-LIGHT^ (mHVEM H86D-L90A double mutein) with mBTLA, mCD160 and mLIGHT as measured by Octet bio-layer interferometry (BLI) technology.

Residues G72 and V74 contribute to the binding interface of the hHVEM:hCD160 and hHVEM:hBTLA complexes (**Fig. 2, F-G and 4 B**), whereas H86 and L90 resides are within the hHVEM:hLIGHT interface in close proximity to hLIGHT Y173, based on the hHVEM:hLIGHT structure (**Fig. 2 H and 4 B**). The mHVEM G72R-V74A double mutation exhibited no binding to mBTLA or mCD160, while it retained wild-type binding to mLIGHT in our cell-cell and cell-protein interaction system (**Fig. 4 A and S3, F-G**). This mHVEM mutant was selected for further analysis and is designated as mHVEM^-BT/160^, denoting loss of BTLA and CD160 binding. The mHVEM H86D-L90A double mutation showed no binding to mLIGHT and wild-type binding to mBTLA and mCD160 (**Fig. 4 A and S3, F-G**). This mHVEM H86D-L90A mutant is thus designated as mHVEM^-LIGHT^, denoting loss of LIGHT binding. Both mHVEM^-BT/160^ and mHVEM^-LIGHT^ proteins were expressed in soluble form and their ligand binding was measured by surface plasmon resonance. The mHVEM^-BT/160^ eliminated binding to both mBTLA/mCD160 while it still retained close to wild-type binding to mLIGHT (**Fig. 4 C**). The mHVEM^-LIGHT^ had approximately 5-fold and 3-fold reduced binding to mBTLA and mCD160, respectively, but had more than a three log-fold decrease in binding to mLIGHT (**Fig. 4 C**).

### mHVEM^-LIGHT^ mice are more susceptible to Yersinia infection

We tested the role of the mouse HVEM muteins, mHVEM^-BT/160^ (G72R-V74A) and mHVEM^-LIGHT^ (H86D-L90A) *in vivo*. We used the CRISPR/Cas9 system to generate two knockin (KI) mouse strains (**Fig. S4 A**). KI homozygous mice having either HVEM mutein were born at the expected frequency with normal size and maturation. Immune cells from homozygous KI mice from either strain expressed a normal surface level of HVEM in different cell types, including splenic CD4^+^ T cells, invariant nature killer T (iNKT) cells, and innate lymphoid cells (ILCs) (**Fig. S4 B**).

Previously, using conditional HVEM knockouts, we reported that HVEM signals in ILC3 are critical for host defense against oral infection with *Yersinia enterocolitica* (*Y. enterocolitica*) (Seo et al., 2018). Importantly, the evidence from whole body LIGHT- deficient mice suggested that this HVEM-mediated protection was dependent on LIGHT, not on BTLA or CD160. These data did not exclude a contribution by other aspects of this network. For example, LT*β*R deficient mice were not tested and LIGHT-LT*β*R interactions are also eliminated when the gene encoding LIGHT is deleted. To test the *in vivo* function of the HVEM muteins, mHVEM^-BT/160^ and mHVEM^-LIGHT^ mice were orally infected with *Y. enterocolitica*. Homozygous mHVEM^-LIGHT^ (KI/KI) mice displayed lower survival, more pronounced weight loss, and large areas of necrosis in the liver and spleen compared with control WT mice (**Fig. 5, A-C**). This severe disease outcome is similar to that observed in *Light* knockout mice (Seo et al., 2018), indicating LIGHT-LT*β*R interactions do not contribute to resistance or cannot overcome the effect of loss of LIGHT binding to HVEM expressed by ILC3. Interestingly, heterozygous mHVEM^-LIGHT^ (KI/+) mice had an intermediate phenotype, with weight loss similar to homozygous mHVEM^-LIGHT^ mice, but they showed better survival than mHVEM^-LIGHT^ mice, as well as reduced necrotic areas and decreased bacterial foci in spleen and liver. Considering that LIGHT binding induces a trimerization of HVEM that likely enhances signaling, an intermediate phenotype might be expected in KI/+ heterozygous mice that would form fewer WT (wild-type) HVEM trimers. In a separate group of *Y. enterocolitica* infections carried out with mHVEM^-BT/160^ mice, animals homozygous for a gene encoding the HVEM mutein that does not bind either IgSF ligand responded similarly to WT mice (**Fig. 5, D-F**). Because of normal experimental variability in bacterial cultures, there was increased weight loss and decreased survival in the WT mice in the series of experiments with mHVEM^-BT/160^ mice (**Fig. 5, D-F**) compared to WT controls in experiments with mHVEM^-LIGHT^ mice (**Fig. 5, A-C**). The key comparison, however, is mHVEM mutein expressing to WT mice within an experiment, and only mHVEM^-LIGHT^ showed a difference. Also, note that clearance was greatly diminished at day 12 only in mHVEM^-LIGHT^ mice and the recovery from weight loss was complete at the end of the experiment in surviving mHVEM^-BT/160^ mice, similar to the WT controls. Therefore, our data suggest that indeed LIGHT is the unique ligand for HVEM in protection from *Y. enterocolitica* and that LIGHT binding to the LT*β*R is not relevant in this context.

**Figure 5.**
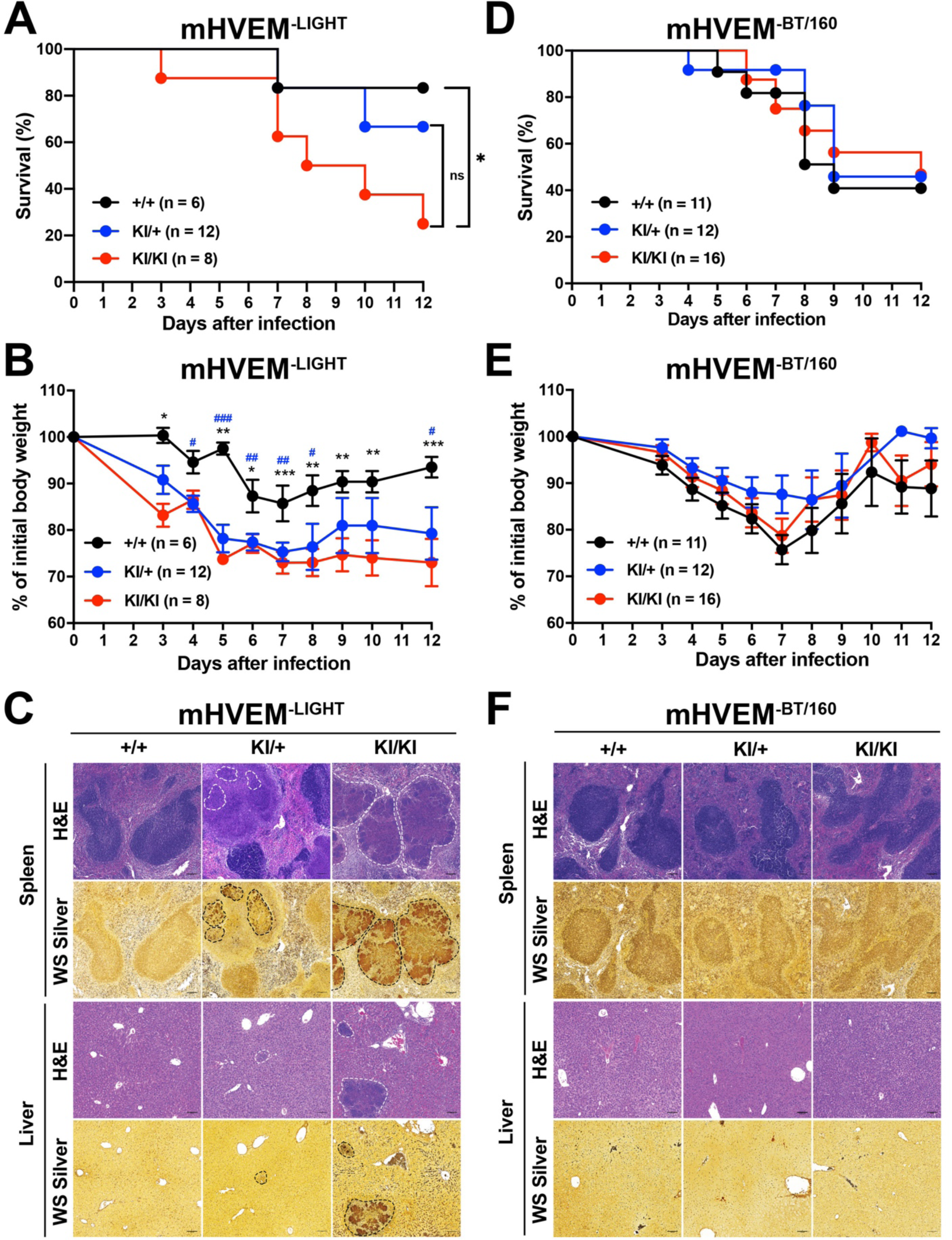
mHVEM^-LIGHT^ mice were more susceptible to *Y. enterocolitica* infection. Male mice were infected with 1.0 x 10^8^ *Y. enterocolitica*. KI = gene knockin. **(A and D)** Survival curves. NS, not significant. \**P* = 0.047 for Log-rank test. **(B and E)** Changes in body weight (% of baseline). \**P* < 0.05; \*\**P* < 0.01; \*\*\**P* < 0.001 (+/+ vs KI/KI) or ^#^*P* < 0.05; ^##^*P* < 0.01; ^###^*P* < 0.001 (+/+ vs KI/+) for two-way ANOVA with Bonferroni’s multiple hypothesis correction. **(C and F)** Representative hematoxylin and eosin (H&E) staining to detect necrotic areas and Warthin-Starry (WS) silver staining to detect bacteria in splenic and hepatic sections from the indicated mice at 7 days after infection. Scale bars, 100 μm. White dotted lines indicate necrotic areas and black dotted lines indicate *Y. enterocolitica*. Data shown are mean ± s.e.m., and represent pooled results from at least two independent experiments having at least three mice per group in each experiment (n= 6-12 mice per group; co-housed littermates). Because of the number of mice that could be handled, experimental data in A-C were done at a different time with a different bacterial culture from D-E.

### mHVEM^-BT/160^ mice are more susceptible to hepatic inflammation

Previous studies have reported that *Btla^-/-^* or *Cd160^-/-^* mice are more susceptible to hepatic injury induced by Concanavalin A (ConA) or by the synthetic glycosphingolipid alpha-galactosylceramide (*α*GalCer) (Iwata et al., 2010; Kim et al., 2019; Miller et al., 2009). We focused on *α*GalCer because of its well-defined mechanism of action as a specific activator of iNKT cells, which are very abundant in intrahepatic lymphocyte populations. When mice are injected with *α*GalCer, iNKT cells are rapidly stimulated and produced many types of pro-inflammatory cytokines, including TNF, IFN*γ*, and IL-4, driving liver injury (Biburger and Tiegs, 2005; Wang et al., 2013). Furthermore, both BTLA and CD160 are expressed by iNKT cells and both molecules served to attenuate production of inflammatory cytokines by iNKT cells during *α*GalCer-induced acute hepatitis (Kim et al., 2019; Miller et al., 2009), providing an example in which two HVEM binding IgSF molecules are required in one cell type. The function of LIGHT in this model has not been reported.

*α*GalCer was injected into female mHVEM^-LIGHT^ and mHVEM^-BT/160^ mice and controls. mHVEM^-LIGHT^ mice presented with a similar phenotype to controls, which at this dose induced only limited *α*GalCer-triggered liver damage and serum ALT activity (**Fig. 6, A-C**). By contrast, larger white spots on the surface of liver and massive hepatic necrotic regions developed in mHVEM^-BT/160^ mice (**Fig. 6, A and B**). Consistently, serum alanine aminotransferase (ALT) activity was elevated in mHVEM^-BT/160^ mice compared with littermate control or heterozygous (KI/+) mice (**Fig. 6 D**). Heterozygous mHVEM^-BT/160^ mice showed an intermediate phenotype, particularly with regard to the ALT measurement. Considering that the IgSF ligand-HVEM interaction is monomeric, this phenotype could reflect HVEM gene haploinsufficiency. These findings suggest that HVEM:BTLA and/or HVEM:CD160 engagement generated negative signaling in iNKT cells, thereby preventing severe *α*GalCer-induced liver injury and hepatitis.

**Figure 6.**
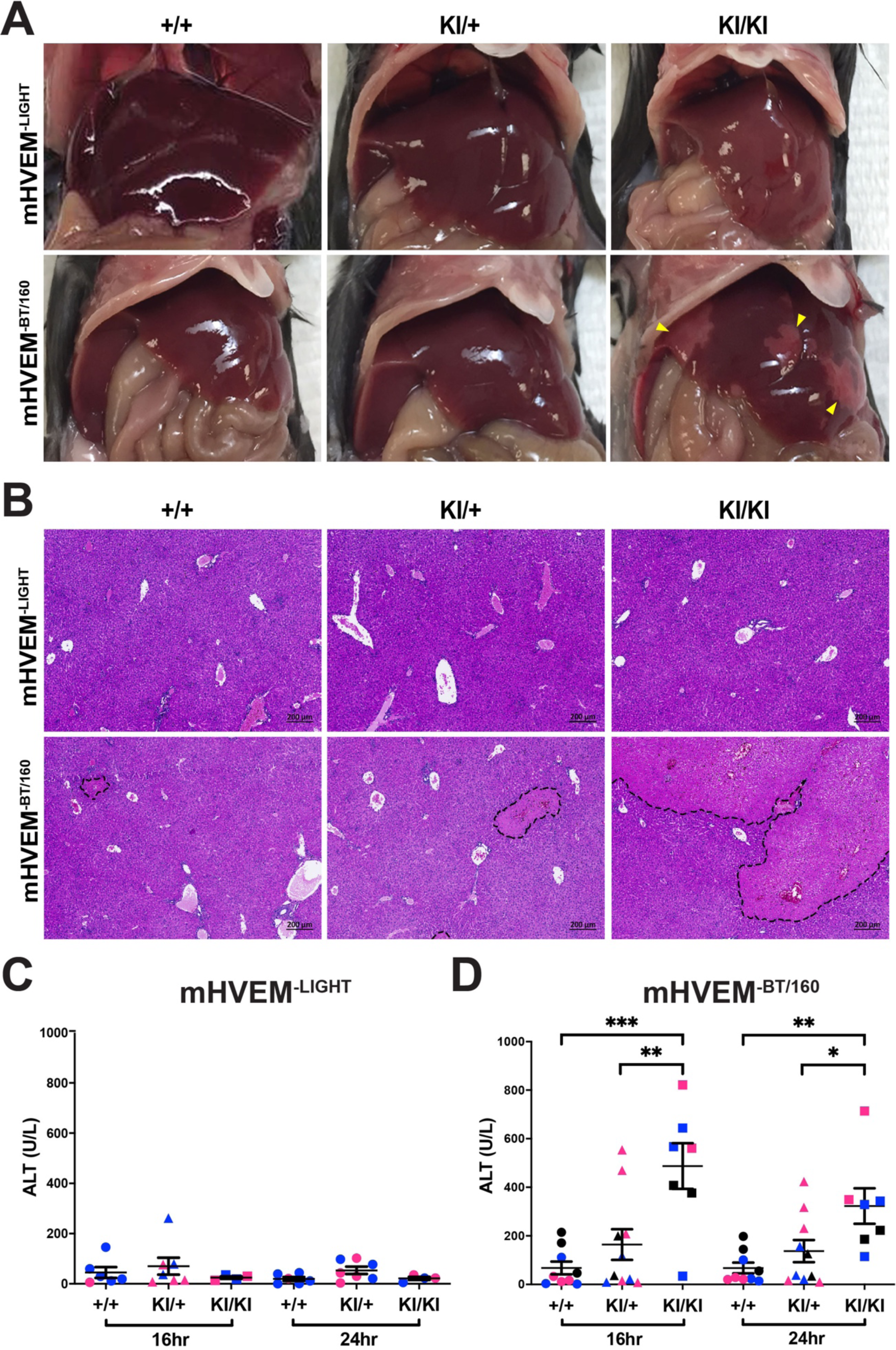
Susceptibility to αGalCer-induced liver injury in mHVEM^-BT/160^ mice. Mice were injected with 2 μg αGalCer by the retro-orbital route. **(A)** Representative images of the liver 24 h after injection. Yellow triangles indicate necrotic areas. **(B)** Representative H&E staining of hepatic sections from the indicated mice 24 h after injection. Black dotted lines indicate the necrotic areas. Scale bars, 200 μm. **(C and D)** Serum ALT activity at 16 and 24 h from the indicated mice. Data shown are mean ± s.e.m.. \**P* < 0.05; \*\**P* < 0.01; \*\*\**P* < 0.001 for one-way ANOVA. Data represent pooled results from at least two independent experiments; each experiment labeled with different colored symbols (n= 4- 10 mice per group; co-housed littermates).

## Discussion

HVEM and its ligands constitute an interacting network of cell surface proteins that affect many aspects of lymphocyte function, as well as the responses of numerous other cells types including eosinophils, keratinocytes, epithelial cells and macrophages in the brain (Doherty et al., 2011; Herro et al., 2018; Shui et al., 2012; Zhu et al., 2016). In order to understand how HVEM functions *in vivo* in this network, and to develop therapeutics based on its mechanisms of action, one important tool is new mouse strains including those that delete HVEM expression in certain cell types (Mintz et al., 2019; Seo et al., 2018), mutants that separate HVEM ligand function from HVEM signaling, and expression of HVEM mutants with selective binding to only certain ligands. Here we report the structures of human orthologs of members this network, including the ternary hHVEM:hLIGHT:hCD160 and binary hHVEM:hLIGHT complexes; we also report the structure of mHVEM in isolation. These structures guided mutagenesis studies that identified HVEM muteins with selective ligand binding. Additionally, we have tested these HVEM muteins in vivo in mouse strains. In this way, without eliminating expression of any member of the network, we provide data indicating that that selective HVEM-ligand interactions are responsible for host defense from enteric bacterial infection and for the prevention of liver inflammation.

In contrast to the homotrimeric structure of LIGHT, BTLA and CD160 proteins are monomers (Compaan et al., 2005; Zhu et al., 2016). Crystallographic and biochemical studies illustrated that hHVEM:hBTLA and hHVEM:hCD160 complexes are characterized by a 1:1 stoichiometry (**Fig. 2, C-D**) (Compaan et al., 2005). Unlike trimeric LIGHT, which directly drives formation of assemblies containing multiple HVEM molecules, monomeric BTLA and CD160 may activate HVEM receptor to promote NF-*κ*B signaling and cell survival (Cheung et al., 2009a; Cheung et al., 2009b) through other mechanisms. The membrane-anchored forms of BTLA and CD160 could drive the localized enrichment of HVEM at cell-cell interfaces, and as a consequence enhance the local concentration of HVEM cytoplasmic domains and associated signaling molecules. Additionally, soluble trimeric LIGHT could contribute by driving the formation of assemblies that bring up to three molecules of HVEM into close proximity, which may facilitate increased local density of HVEM:BTLA and HVEM:CD160 complexes. The recognition interfaces in the ternary hHVEM:hLIGHT:hCD160 complex are similar to those in the binary hHVEM:hCD160 and hHVEM:hLIGHT complexes, suggesting that little molecular accommodation is required for HVEM to simultaneously engage two types of binding partners. It remains to be determined under which conditions HVEM concurrently binds LIGHT and one of its IgSF ligands, if a trimeric HVEM:LIGHT complex can contain mixed IgSF binding partners, both CD160 and BTLA, and importantly, whether these interactions enhance BTLA- or CD160- mediated signals. Furthermore, LIGHT can be expressed in membrane bound or soluble forms, and it is not known if the membrane-bound form also can bind HVEM simultaneously with BTLA or CD160. Previously, it was suggested that when LIGHT and BTLA are presented on the same cell membrane, membrane LIGHT might limit BTLA binding in trans due to steric incompatibilities associated with the position of the LIGHT and IgSF binding sites on HVEM relative to the cell membrane (Steinberg et al., 2011). In humans, the stalk region of LIGHT is 35 amino acids, while for BTLA it is only 24 amino acids. For hCD160, it is 17 amino acids for the glycosylphosphatidylinositol (GPI)-linked form and 19 amino acids for the transmembrane form. These constraints would position BTLA and CD160 too close to the cell membrane to bind HVEM together with LIGHT (**Fig. S5**). Therefore, it is possible that the membrane bound and secreted forms of LIGHT could have different impacts on HVEM:BTLA and HVEM:CD160 binding, based on their position relative to the cell membrane, but additional *in vitro* and *in vivo* studies will be required to verify this.

Whole body and cell-type-specific gene knock outs have provided important insights into the function of HVEM and its binding partners (Mintz et al., 2019; Seo et al., 2018). Elimination of expression of one member of this network, however, could have complex effects on others. For example, deletion of LIGHT not only eliminates LIGHT-HVEM interaction, but also the LIGHT-LT*β*R interaction. It is also possible that LIGHT deletion might provide more LT*β*R available for binding to LT*αβ*2, and in humans, blockade of LIGHT may alter the degree of inhibition of TL1A and FasL by DcR3, a decoy receptor not present in mice. Although analysis of no single mutation can discriminate between all these possibilities, we set out to test the importance *in vivo* of pairwise interactions in the HVEM network in a context in which expression of all of the proteins was maintained. To do this, we mutated solvent accessible amino acids in mHVEM that are close to the ligand binding interfaces defined by structural analyses. We succeeded in identifying mHVEM muteins with selective binding *in vitro* for either LIGHT or for the two IgSF ligands. These HVEM proteins were expressed at normal amounts on cells in genetically altered mouse strains and were tested *in vivo* following oral infection with *Y. enterocolitica* and following injection with *α*GalCer to activate iNKT cells to cause liver inflammation. These data demonstrate a high degree of ligand selectivity in this more complete network. Our data show that LIGHT-HVEM interactions are required for host defense against *Y. enterocolitica*. In mice that retain normal expression of LIGHT and HVEM, but in which only the ability of these proteins to interact was greatly diminished, bacteria spread and weight loss were increased and survival was diminished. The phenotype was similar to mice deficient for HVEM in T cells and ILC3, or in whole body knockout mice lacking LIGHT expression. There was no effect on the host response in mice in which HVEM binding to CD160 and BTLA was diminished. Similarly, liver inflammation was dependent on CD160 and/or BTLA interacting with HVEM. As suggested by other studies (Iwata et al., 2010; Kim et al., 2019; Miller et al., 2009), this behavior may be due to the loss of inhibitory signaling in the iNKT cells that initiate this inflammatory response. It was not greatly dependent on LIGHT binding to HVEM, suggesting LIGHT induced HVEM trimerization is not a major factor in promoting or inhibiting BTLA and CD160 signaling in this system.

It is not known why individual HVEM ligands are important for mediating biologic effects in particular contexts, and how the great difference in binding affinity between LIGHT and the IgSF binding partners contribute to these processes. All of the ligands activate NF-*κ*B proteins (Cheung et al., 2009b), and there is no evidence that they employ different mechanisms for signaling through HVEM. Tissue context is likely critical in some cases. For example, it is not surprising that intestine epithelial HVEM interacts mainly with CD160 expressed by intraepithelial lymphocytes (IEL), because these cells are in continual contact with the epithelium (Shui et al., 2012), and CD160 is the only HVEM binding partner IEL highly express. Reverse signaling by HVEM through either CD160 or BTLA could drive the biology in other instances, as reported recently for the germinal center response (Mintz et al., 2019) or in the liver inflammation model (Iwata et al., 2010; Kim et al., 2019; Miller et al., 2009). Ultimately, a deeper understanding of the biologic effects of HVEM may permit the safer use of muteins and other reagents in a therapeutic context; for example, in cancer immunotherapy, where soluble HVEM has shown benefit in a mouse model of lymphoma (Pasero and Olive, 2013; Sedy and Ramezani-Rad, 2019) or for treating inflammatory diseases.

## Materials and methods

### Molecular Cloning and Mutagenesis

A portion of the hHVEM gene encoding residues L39-C162 and mHVEM encoding residues Q39-T142 were amplified by PCR and the resulting DNA fragments were digested with endonucleases BglII and AgeI and ligated into plasmid pMT/BiP/V5-His for His-tag fusion protein production in *Drosophila* S2 cells. DNA fragment encoding the amino acid sequence “HHHHHHG” fused to hLIGHT (L83-V240) was cloned into pMT/BiP/V5-His. The mCD160 gene encoding residues 30I-154H with the C-terminus fused with amino acids “HHHHHHGGGGSGLNDIFEAQKIEWHE” was cloned into pET3a. The DNA sequences encoding a protein biologic composed of mHVEM residues (Q39- Q206) followed by human IgG1 and a subsequent hexa-His tag sequences were cloned into pcDNA 3.3 vector (Life technologies) using In-fusion HD cloning enzyme premix (Clontech). DNA fragment encoding the amino acid sequence “HHHHHHGG” fused to the N-terminus of the single chain homotrimeric mLIGHT extracellular domain (G73-V239) connecting by two (GGGGS)4 linkers was cloned into pcDNA 3.3 vector (Life technologies).

A DNA fragment encoding residues of L39-V202 of hHVEM was cloned into an engineered pGFP-N1 vector (Clontech) for expression as a protein fused with a PD-L1 trans-membrane domain followed by the fluorophore eGFP at the C-terminus. The hHVEM mutant library was generated using the QuickChange II Site-Directed Mutagenesis Kit (Agilent Technologies). Full length of WT mHVEM and mutants were cloned into pmCherry-N1 vector (Clontech), respectively. Full length of mBTLA was cloned into pEGFP-N1 vector (Clontech). Full length of mLIGHT was cloned into pIRES2-EGFP vector (Clontech), which contains a subsequent IRES (Internal Ribosome Entry Site) sequence following by a fluorescent EGFP ORF.

### Protein Production and Purification

All hHVEM, hLIGHT and mHVEM proteins were expressed and purified as previously described (Liu et al., 2015). The extracellular domains of hHVEM (L39-C162), hLIGHT (L83-V240) and mHVEM (Q39-T142) were separately cloned into the pMT/BiP/V5-His A vector (Invitrogen) and co-transfected into *Drosophila* S2 cells with the pCoBlast (Invitrogen) plasmid at a 20:1 ratio. A stable cell line was selected with Blasticidin following the manufacture’s protocol (Invitrogen). All hHVEM, hLIGHT and mHVEM expression were induced with copper sulfate (500 μM final concentration). The proteins from filtered culture supernatants were purified by Ni-NTA column (QIAGEN) and size exclusion chromatography (HiLoad Superdex 75; Amersham). The single chain hCD160- hHVEM fusion protein was expressed in *Drosophila* S2 cells and purified to homogeneity as previously described (Liu et al., 2019). The mCD160 protein was purified as inclusion bodies and refolded as previously described (Liu et al., 2019). The expression vectors encoding mHVEM (Q39-Q206) fused with human IgG1 and a subsequent hexa-His tag sequences were transfected into Expi293 (Gibco) cells using ExpiFectamine 293 transfection kit (Gibco) and the resulting proteins were purified using Ni-resins (Qiagen). The vector encoding a hexa-His tag fused to a single chain homotrimeric mLIGHT extracellular domain (G73-V239) connecting by two (GGGGS)4 linkers was transfected into Expi293 (Gibco) cells using the ExpiFectamine 293 transfection kit (Gibco) and the resulting proteins were purified using Ni-resins (Qiagen) and size exclusion chromatography (HiLoad Superdex 75; Amersham). The resulting purified mLIGHT proteins were used freshly.

### Cell culture

Transformed *E. coli* cells were cultured in LB (Lysogeny Broth) medium supplemented with 100 mg/L Carbenicillin at 37 °C. Transfected *Drosophila* S2 cells were cultured in complete Schneider’s *Drosophila* medium (Life Technologies) supplemented with 10% heat-inactivated fetal bovine serum in the presence of 25 mg/L Blasticidin for establishing stable cell lines. Protein expression in *Drosophila* S2 cell lines was induced in Express Five SFM medium (Life Technologies) in the presence of 500mM CuSO4 at 25 °C. Expi293 cells were maintained in DMEM (Corning) with 10% FBS at 37 °C with 5% CO2. The transfected Expi293 cells were cultured at 37 °C with 5% CO2 for flow cytometry analysis or at 30 °C with 5% CO2 for protein expression.

### Crystallization, Structure Determination and Refinement

The purified hHVEM and hLIGHT proteins were concentrated separately and mixed in a 1:1 molar ratio to generate the hHVEM:hLIGHT complex, at a concentration of 3 mg/mL in 10 mM HEPES, pH 7.0 and 150 mM NaCl solution. The resulting hHVEM:hLIGHT complex was crystallized by sitting drop vapor diffusion using 0.5 μL of protein and 0.5 μL of precipitant composed of 0.1 M Bis-Tris, pH5.5, 0.2 M MgCl2 and 9% PEG3350. Crystals were cryo-protected by immersion in crystallization buffer supplemented with 20% of glycerol, and flash-cooled in liquid nitrogen. The purified single chain hCD160-hHVEM proteins and hLIGHT were concentrated separately and mixed in a 1:1 molar ratio to generate the hHVEM:hLIGHT:hCD160 complex at a concentration of 5 mg/mL in 10 mM HEPES, pH 7.0 and 150 mM NaCl solution. The resulting hHVEM:hLIGHT:hCD160 complex was crystallized by sitting drop vapor diffusion using 0.5 μL of protein and 0.5 μL of precipitant composed of 12% (W/V) PEG3350 and 4% (V/V) tacsimate. Crystals were cryo-protected by immersion in crystallization buffer supplemented with 20% ethylene glycerol, and flash-cooled in liquid nitrogen. The purified mHVEM was concentrated to 3 mg/mL in 10 mM HEPES, pH 7.0 and 150 mM NaCl solution and then crystallized by sitting drop vapor diffusion using 0.5 μL of protein and 0.5 μL of precipitant composed of 90% (V/V) solution A with 0.2 M lithium sulfate, 0.1 M sodium acetate/acetic acid, pH4.5, 30% (W/V) PEG 8000 and 10% (V/V) solution B with NDSB-211. Crystals were cryo-protected by immersion in crystallization buffer supplemented with 40% of glycerol, and flash-cooled in liquid nitrogen.

Diffraction data from the hHVEM:hLIGHT complex were collected at Brookhaven National Laboratory (BNL) beamline X29 (**Table 1**). Diffraction data from hHVEM:hLIGHT:hCD160 complex and mHVEM were collected at Advanced Photon Source Sector 31, Argonne National Laboratory (**Table 1**). All diffraction data were integrated and scaled with HKL2000 (Otwinowski and Minor, 1997). Phases of the hHVEM:hLIGHT complex were calculated by molecular replacement using the existing PDB structures 4KG8 and 4FHQ as the starting models and the software Molrep in the CCP4 package (Winn et al., 2011). Phases of hHVEM:hLIGHT:hCD160 complex were calculated by molecular replacement using the existing PDB structure 6NG9 and hHVEM:hLIGHT complex (PDB entry 4RSU) as the starting models and the software Molrep in the CCP4 package (Winn et al., 2011). Phases of mHVEM were calculated by molecular replacement using the existing PDB structure 4FHQ as the starting model and the software Molrep in the CCP4 package (Winn et al., 2011). Electron density maps were manually inspected and improved using COOT (Emsley et al., 2010). Following several cycles of manual building in COOT and refinement in REFMAC5, the hHVEM:hLIGHT complex Rwork and Rfree converged to 18.4% and 22.6%, respectively (Emsley et al., 2010; Winn et al., 2011).

### Mutagenesis screening by flow cytometry binding assays

500 ng wild type and mutants of hHVEM-GFP fusion plasmids in 50 μL PBS were mixed with 50 μL of 0.04 M polyethylenimine (PEI), respectively. The mixtures were kept still for 10 min and then added separately to a 24-well plate with each well containing 1mL of 10^6^/mL HEK293-Freestyle cells (Invitrogen). The transfected cells were cultured by shaking at a speed of 200 rpm at 37 °C for 72 h followed the transfection, and then the cells were collected and resuspended in PBS. Cells from each well were further diluted to 10^6^ cells/mL.

100 μL of the diluted transfected cells were incubated separately with hCD160- 6×His tag, hBTLA-6×His tag (R&D systems) and hLIGHT-6×His tag proteins (made by the methods described above) in the mixtures with anti-6×His tag PE-labeled antibody (Abcam) for 20 min on ice. The cells were subsequently spun down, washed once and resuspended in 100 μL PBS buffer containing additional 0.5% BSA and then subjected to flow cytometric analysis. The cells were gated on GFP positive cells to ensure hHVEM expression and analyzed for the percentage of PE positive cells. The binding of wild-type hHVEM was normalized as 1. The relative binding of hHVEM mutants were calculated by comparing the PE positive cell percentage to the control wild-type hHVEM groups. The error bars reflect the results of three independent experiments.

The mHVEM, mBTLA and mLIGHT constructs were transfected into HEK293 FreeStyle (Life technologies) cells using PEI (Linear Polyethylenimine with molecular weight of 25000; Polysciences Inc.). After 2∼3 days, the cells were harvested and diluted to 10^6^/mL. For measuring cell-cell interactions, 100 μL of cells expressing mHVEM- mCherry proteins were mixed with 100 μL of cells expressing mBTLA-EGFP or mLIGHT- IRES-EGFP proteins and then subjected to shaking (900 RPM) at room temperature for 2 h. These cells were further recorded and analyzed by flow cytometry. For protein staining, 100 μL of cells expressing mHVEM-mCherry proteins were mixed with 0.3 μg mBTLA- penta-His-tag/mCD160-biotin proteins and 0.5 μg of green fluorescent anti-His-tag (Abcam; Cat: ab1206)/Alexa Fluor 488 conjugated streptavidin (Life technologies; Cat: S11223) proteins. The cells were incubated for 30 min with shaking at room temperature and washed once by PBS containing 0.2% BSA (PBS-BSA). The cells were re-suspended in 100 μL of PBS-BSA and analyzed by flow cytometry.

### Measuring affinities of mHVEM muteins using Octet bio-layer interometry (BLI) technology

For measuring binding affinities, mHVEM-hIgG1 was immobilized on the sensors (ForteBio) and then challenged with different concentrations of mLIGHT, mBTLA or mCD160. The results were exported and then analyzed using Prism 5 (GraphPad Software). Final response curves were generated after subtracting the responses of the control groups. The equilibrium dissociation constants (KD) of the mHVEM-hIgG1 interaction with mLIGHT were calculated based on the response curves by fitting the data to the equation Y=Bmax X / (X + KD) (Y is the averaged maximum response of each experiments. X is the concentration of the analytes and Bmax is the maximum specific binding. The equilibrium dissociation constants (KD) of mHVEM-hIgG1 interaction with mBTLA or mCD160 were calculated based on the 1:1 Langmuir model.

### Generation of mHVEM mutant mice

The mHVEM mutant mice were generated using the CRISPR/Cas9 system. The transgenic mouse core of the UC San Diego Moores Cancer Center injected the sgRNA- Cas9 complex plus a specific single-stranded DNA (ssDNA) homology directed repair (HDR) template into C57BL/6 pronuclear embryos. All materials of the CRISPR/Cas9 system were designed and ordered from Integrated DNA Technologies (IDT, Newark, NJ). Two specific sgRNAs targeted exon 3 of the *Tnfrsf14* locus, sgRNA-1 for *Tnfrsf14*^G72R/V74A^ (mHVEM^-BT/160^): 5’-CAGGTCTGCAGTGAGCATAC-3’ and sgRNA-2 for *Tnfrsf14*^H86D/L90A^ (mHVEM^-LIGHT^): 5’-ACATATACCGCCCATGCAAA-3’. Two specific ssDNAs were used as HDR templates, mHVEM^-BT/160^: 5’- TGGCTGCAGGTTACCATGTGAAGCAGGTCTGCAGTGAGCACACGCGTACAGCGTGTGCCCCCTGTCCCCCACAGACATATACCGCCCATGCA-3’ and mHVEM^-LIGHT^: 5’- CAGGCACAGTGTGTGCCCCCTGTCCCCCACAGACATATACAGCGGACGCTAATGGCGCTAGCAAGTGTCTGCCCTGCGGAGTCTGTGATCCAGGTAGGA-3’. For screening, we created a new restriction enzyme site near the PAM sequence, which did not alter the amino acid sequence. A new MluI or NheI site was thereby created in the knockin genomes of the mHVEM^-BT/160^ or mHVEM^-LIGHT^ mice, respectively. The F0 founder pups were screened for exon 3 of the *Tnfrsf14* locus by enzyme digestion and PCR using the primers Hvem-exon3-F1 (5’-GTACAGTGTTCAGTTCAGGGATAG-3’) and Hvem-exon3- R1 (5’-AGCAGGAAAGAACCTCTCATTAC-3’). The *Tnfrsf14* exon 3 sequences were cloned and sequenced from each line of founder mice that had undergone HDR repair. The successfully HDR repair F0 founders were first backcrossed to the WT C57BL/6 strain. Germ-line transmission of each line of mHVEM mutant mice (N1) was verified by PCR and restriction enzyme digestion analysis. Testing for potential off-target genes, analyzed by the software from IDT, and homologous sequences were confirmed by PCR using a specific pair of primers on each gene and sequencing at the N1 generation. We examined six potential off-target genes from mHVEM^-BT/160^ strain and four genes from mHVEM^-LIGHT^ strain. Two and four founders from mHVEM^-BT/160^ or mHVEM^-LIGHT^ strain, respectively, were verified and backcrossed again to the WT C57BL/6 mice. After two backcrosses with C57BL/6 mice, we obtained heterozygous (KI/+) mice (N2) from each mHVEM mutant strain. We obtained homozygous offspring (N2F1) by intercrossing the N2 generation of KI/+ mice. Age and gender matched cohoused littermates were used for experiments. All mice were bred and housed under specific pathogen-free (SPF) conditions in the vivarium of La Jolla Institute for Immunology (LJI) and all animal experimental procedures were approved by the LJI Animal Care and Use Committee.

### Bacterial infection

*Yersinia enterocolitica* strain WA-C (pYV::CM) was prepared as described previously (Seo et al., 2018; Trulzsch et al., 2004). Briefly, *Yersinia* were grown overnight in LB broth at 30°C, and the overnight culture was expanded with fresh medium for 6 h. Bacteria were washed and diluted with PBS. Co-housed male littermates were infected by oral gavage with 1×10^8^ c.f.u. of *Y. enterocolitica*. Infected mice were analyzed by measurement of body weight daily and tissues were harvested at 7 days after infection for determination of bacterial c.f.u. and histologic analysis as described previously (Seo et al., 2018).

### Hepatic inflammation

Co-housed female littermates were inoculated with 2 μg *α*GalCer (KRN7000, Kyowa Kirin Research, La Jolla, California) in a total volume of 200 μl PBS by retro-orbital injection. Serum ALT activity was measured using a colorimetric/fluorometric assay kit (K752, Biovision) at 16 or 24 h after injection. Hepatic tissues were collected and the necrotic areas were determined using H&E staining at 24 h after *α*GalCer treatment.

### Statistics analysis

All data were randomly collected and analyzed using Microsoft Office Excel and GraphPad Prism 8 software. Data were shown as mean with the standard error of the mean (s.e.m.). The detail of statistical analysis methods and the representing number of mice (n) is indicated in each figure legend. Statistical significance is indicated by * *P* < 0.05; \*\**P* < 0.01; \*\*\**P* < 0.001.

### Online supplemental material

Fig. S1 illustrates the network of interactions between HVEM and its binding partners. Fig. S2 shows the binding interface between hHVEM and hLIGHT. Fig. S3 shows the relative binding affinities of the HVEM mutants with BTLA, CD160, and LIGHT. Fig. S4 shows the outcome of CRISPR-Cas9 editing of exon 3 of the *Tnfrsf14* locus and also that mHVEM^-BT/160^ and mHVEM^-LIGHT^ mice have normal surface HVEM expression. Fig. S5 illustrates a model for the stalk regions of BTLA, CD160, and LIGHT. Table 1 shows data collection and refinement statistics of the crystal structures of the hHVEM:hLIGHT, hHVEM:hLIGHT:hCD160, and mHVEM complexes and proteins.

## Acknowledgments

We thank the staff of X29A beam lines at the National Synchrotron Light Source. Use of the National Synchrotron Light Source, Brookhaven National Laboratory, was supported by the U.S. Department of Energy, Office of Science, Office of Basic Energy Sciences, under Contract No. DE-AC02-98CH10886. Data for parts of this study were collected at beamline X29A of the National Synchrotron Light Source. Financial support comes principally from the Offices of Biological and Environmental Research and of Basic Energy Sciences of the US Department of Energy, and from the National Center for Research Resources (P41RR012408) and the National Institute of General Medical Sciences (P41GM103473) of the National Institutes of Health. Use of the Advanced Photon Source, an Office of Science User Facility operated for the U.S. Department of Energy (DOE) Office of Science by Argonne National Laboratory, was supported by the U.S. DOE under Contract No. DE-AC02-06CH11357. Use of the Lilly Research Laboratories Collaborative Access Team (LRL-CAT) beamline at Sector 31 of the Advanced Photon Source was provided by Eli Lilly Company, which operates the facility. We also acknowledge support from the Albert Einstein Cancer Center (P30CA013330), the Einstein Crystallographic Core X-ray Diffraction Facility supported by NIH Shared Instrumentation Grant S10 OD020068 and the Albert Einstein Macromolecular Therapeutics Development Facility. This work was partially supported by the Price Family Foundation and contributions to the Albert Einstein Center for Experimental Therapeutics by Pamela and Edward S. Pantzer. We thank Jun Zhao and Ella Kothari from transgenic mouse core of the UC San Diego Moores Cancer Center (supported by NIH grant P30CA023100), Zbigniew Mikulski, Angela Denn, and Katarzyna Dobaczewska from LJI Histology Core (supported by NIH grant P30 DK120515 from San Diego Digestive Diseases Research Center), Kristine Suchey, Mindy Hockaday and the staff from LJI animal facility (DLAC) and Cheryl Kim from LJI Flow Cytometry Core (supported by NIH grant S10RR027366). Support for M.K. from NIH grants U01 AI125955 and P01 DK46763 and support for Ting- Fang from the Academia Sinica-UC San Diego Talent Development Program from Academia Sinica, Taiwan, R.O.C.

## Author contributions

W. Liu designed and performed the experiments to determine the structures and screen the mutants. S.C. Garrett-Thomson helped on the mutagenesis screening. E. Fedorov, U.A. Ramagopal and J.B. Bonanno helped on the structural determination. T.-F. Chou, K. Kakugawa, and H. Cheroutre designed and generated knockin mice. T.-F. Chou designed and performed the mouse experiments. G.-Y. Seo helped on the bacterial infection. S.C. Almo and M. Kronenberg conceived, supervised and managed the project. W. Liu, T.-F. Chou, M. Kronenberg, and S.C. Almo wrote the paper.

Disclosures: The authors declare no competing interests exist.

## Supplemental Material

**Figure S1.**
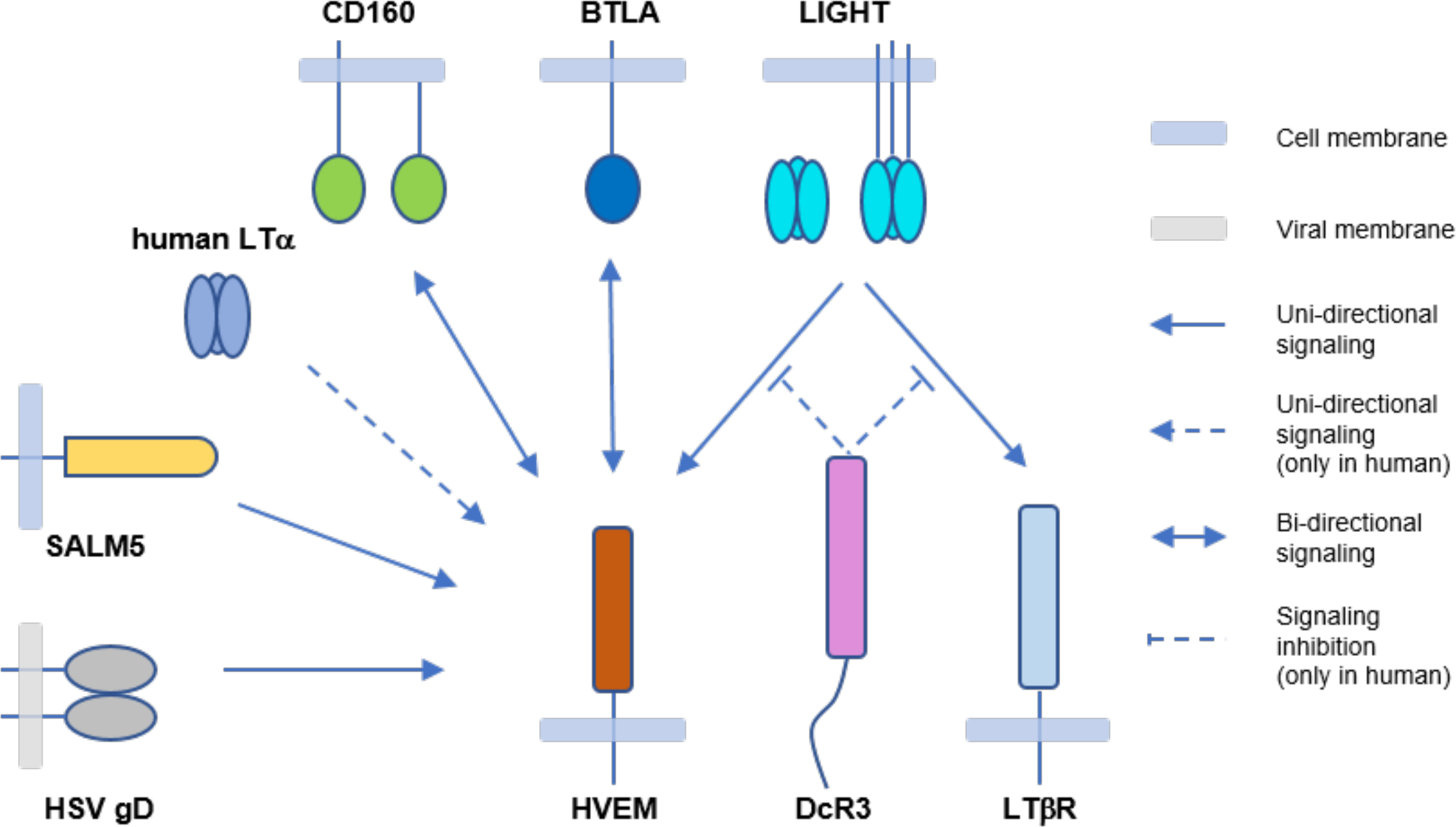
A diagram of the HVEM interaction network. HVEM can be activated by HSV gD, SALM5, CD160, BTLA, LIGHT, and for human HVEM, weakly by LT*α*. The interactions of HVEM with CD160 and BTLA result in bi-directional signaling to activate CD160 and BTLA as well as HVEM. LIGHT also engages LT*β*R besides HVEM, whereas these interactions can be neutralized by soluble DcR3 in humans.

**Figure S2.**
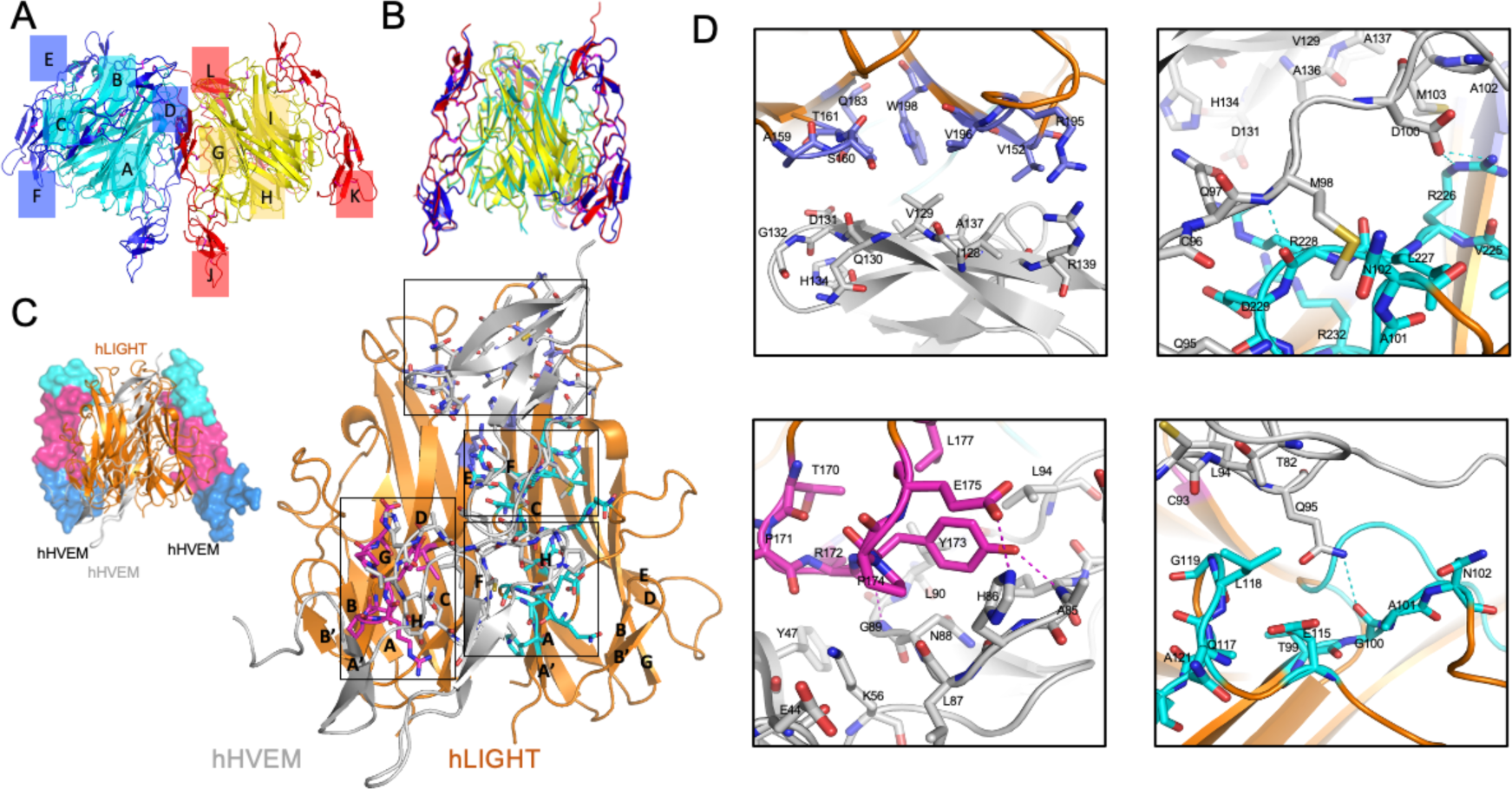
Overall structure of hHVEM:hLIGHT complex and the binding interface between hHVEM and hLIGHT. **(A and B)** One asymmetry unit contains 6 independent chains of hLIGHT (cyan and yellow cartoon) and 6 independent chains of hHVEM (blue and red cartoon) forming two independent 3:3 hHVEM:hLIGHT complexes. Each chain is labeled in the figure. **(A)** Side view of the two hHVEM:hLIGHT complexes in one asymmetry unit. **(B)** Side view of the superimposition result of the two hHVEM:hLIGHT complexes. **(C)** The overall structure of the hHVEM:hLIGHT complex (top left) and magnified view of one copy hHVEM binding to two adjacent hLIGHT monomers (bottom right). The hLIGHT is shown as orange cartoon. The hHVEM is presented as grey cartoon for one copy and CRD colored surface for two copies. **(D)** Magnified views of the binding interface between hHVEM and hLIGHT. The residues from the “upper” region of the hHVEM:hLIGHT complex are shown as marine color sticks on the top left panel. The residues from the AA” and GH loops part of the “lower” region of the hHVEM:hLIGHT interface are shown as cyan sticks on the top right and bottom right panels. The residues from the DE loop part of the “lower” region of the hHVEM:hLIGHT interface are shown as magenta sticks on bottom left panel. The residues of hHVEM contributing to the interface are presented as grey sticks. The interaction interface of the “upper” region between hLIGHT and hHVEM (top left panel). The interaction interface between the GH loop of hLIGHT and hHVEM (top right panel). The interaction interface between the DE loop of hLIGHT and hHVEM (bottom left panel). The interaction interface between the AA’ loop of hLIGHT and hHVEM (bottom right panel).

**Figure S3.**
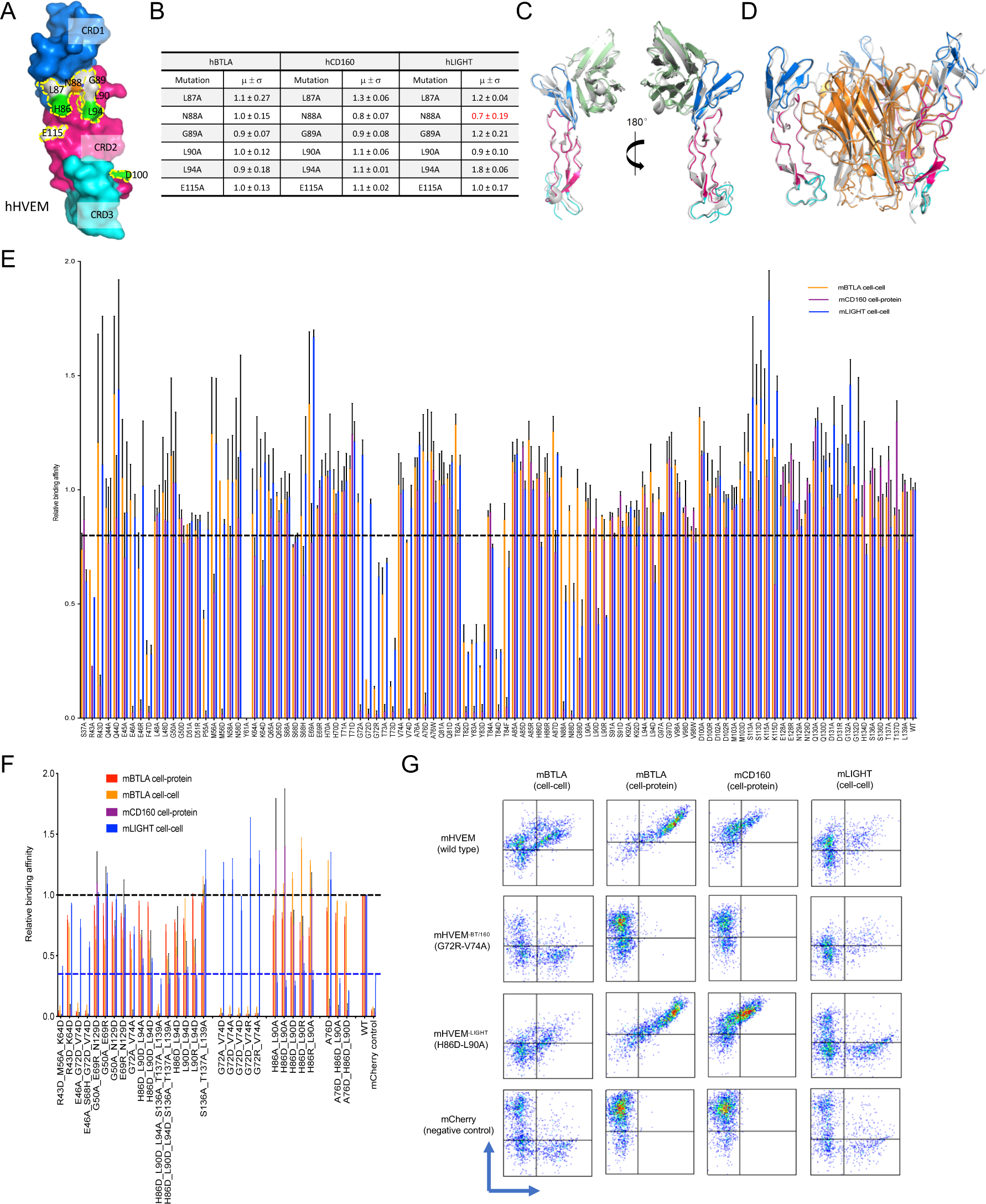
Relative binding affinities of HVEM mutants with BTLA, CD160, and LIGHT. **(A and B)** The hHVEM mutants were expressed on cell surface and were stained by hCD160, hBTLA, and hLIGHT proteins. The relative binding affinities were measured by flow cytometry. Error bars represent results from at least triplicates. **(A)** shows the positions of the hHVEM mutation residues. The residue hLIGHT Y173 (highlighted in yellow) is shown as yellow stick in the structure. **(B)** shows the relative binding affinities of the hHVEM mutants. **(C)** Superimposition of the hHVEM:hCD160 from the ternary complex with hHVEM:hCD160 complex alone (grey cartoon, PDB entry 6NG3). **(D)** Superimposition of the hHVEM:hLIGHT from the ternary complex with hHVEM:hLIGHT complex alone (grey cartoon, PDB entry 4RSU). **(E)** Relative binding affinities of mHVEM single residue muteins with its ligands. Error bars represent results from at least triplicates. **(F)** Relative binding affinities of mHVEM multiple-residue muteins with its ligands. Error bars represent results from at least triplicates. **(G)** Representative flow cytometry results. The vertical axis is the mCherry fluorescence indicating mHVEM-expressing cells and the horizontal axis is the green fluorescence staining of fusion proteins or binding partner- expressing cells, as indicated.

**Figure S4.**
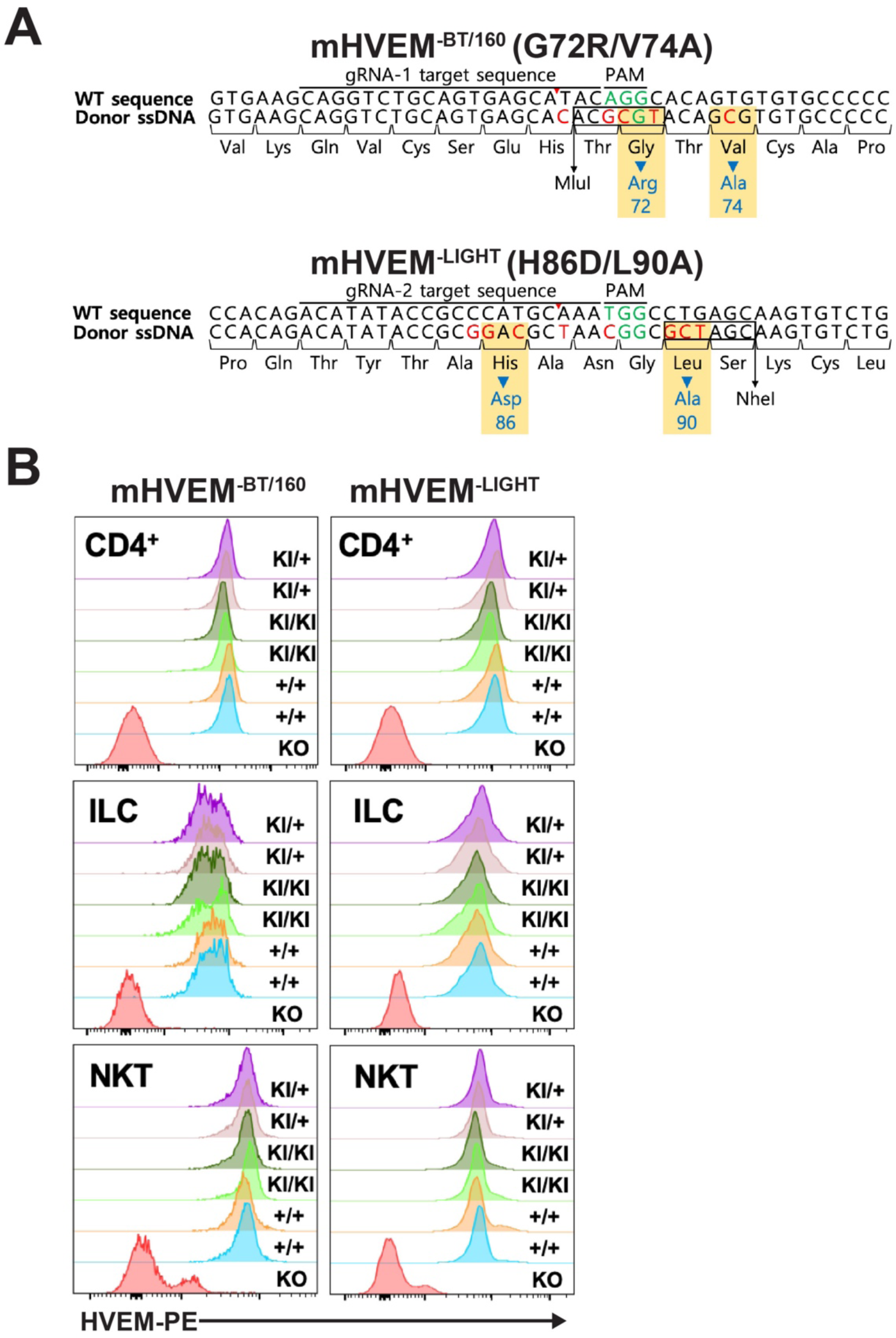
Normal surface HVEM expression in mHVEM mutant mice. **(A)** Schematic of nucleotide sequences of the HVEM gene in mHVEM mutant mouse strains (G72R/V74A: mHVEM^-BT/160^, loss of BTLA and CD160 binding; H86D/L90A: mHVEM^-LIGHT^, loss of LIGHT binding) that were generated by CRISPR-Cas9 editing of exon 3 of the *Tnfrsf14* locus. Red letters indicate mutated nucleotides. Green letters indicate PAM sequence. Blue letters indicate mutated amino acids. Black box shows restriction enzyme sites. **(B)** HVEM surface expression level of splenic CD4^+^ T cells, ILC (CD3^-^Lin^-^CD90.2^+^), and iNKT cells (TCR*β*^+^, CD1d tetramer^+^) from the indicated mice were determined by flow cytometry. HVEM-knockout (KO) mouse plays as a negative control of HVEM staining. KI = knockin allele.

**Figure S5.**
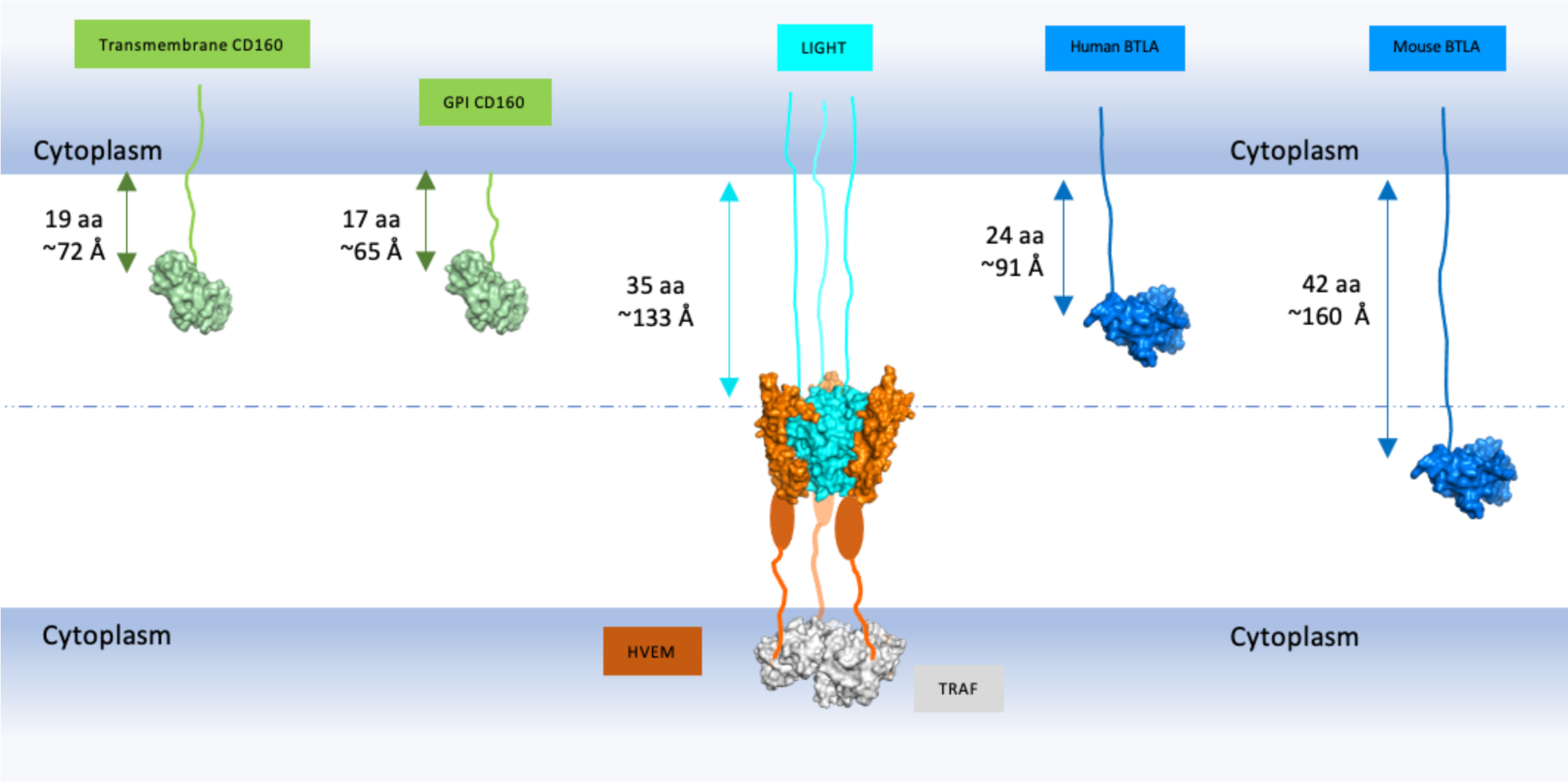
Predicted maximum lengths of LIGHT, CD160 and BTLA stalk regions. The globular domains of LIGHT, CD160 and BTLA are shown as surface structures and colored as cyan, green and blue, respectively. The CRDs of HVEM are shown as orange surfaces and the remainder of the CRD regions that were not visible in the structures are shown as orange ovals. The cytoplasmic TRAF molecule is shown as a grey surface. The stalk regions that connect the extracellular globular domains to the transmembrane segments are shown as lines. The maximum lengths of the stalk regions are calculated as if they adopt the fully extended structures. The length of GPI-anchored CD160 stalk region in the figure does not include the GPI length. Amino acids are denoted as “aa” in the figure. This figure indicates that when human membrane LIGHT binds to HVEM, the longer stalk lengths of LIGHT may prevent BTLA and CD160 binding to HVEM.

## Notes

### Competing Interest Statement

The authors have declared no competing interest.

